# Child Growth Predicts Brain Functional Connectivity and Future Cognitive Outcomes in Urban Bangladeshi Children Exposed to Early Adversities

**DOI:** 10.1101/447722

**Authors:** Wanze Xie, Sarah K.G. Jensen, Mark Wade, Swapna Kumar, Alissa Westerlund, Shahria H. Kakon, Rashidul Haque, William A. Petri, Charles A. Nelson

**Author notes:** Correspondence concerning this article should be addressed to: Charles A. Nelson, Laboratories of Cognitive Neuroscience, Division of Developmental Medicine, Boston Children’s Hospital, 1 Autumn Street, Boston, MA 02215.

## Abstract

**Background:** Faltered growth has been shown to affect 161 million children worldwide and derail cognitive development from early childhood. The neural pathways by which growth faltering in early childhood affects future cognitive outcomes remain unclear, which is partially due to the scarcity of research using both neuroimaging and sensitive behavioral techniques in low-income settings. We employed EEG to examine the association between growth faltering and brain functional connectivity and whether brain functional connectivity mediates the effect of early adversity on cognitive development.

**Methods:** We recruited participants from an urban impoverished neighborhood in Dhaka, Bangladesh. One sample consisted of 85 children whose EEG and growth measures (height for age, weight for age, and weight to height) were collected at 6 months and cognitive outcomes were assessed at 27 months. Another sample consisted of 115 children whose EEG and growth measures were collected at 36 months and IQ scores were assessed at 48 months. Path analysis was used to test the effect of growth measures on cognitive outcomes through brain functional connectivity.

**Findings:** Faltered growth was found to be accompanied by overall increased functional connectivity in the theta and low-beta frequency bands for the 36-month-old cohort. For both cohorts, brain functional connectivity was negatively predictive of later cognitive outcomes at 27 and 48 months, respectively. Faltered growth was found to have a negative impact on children’s IQ scores in the older cohort, and this effect was found to be mediated by brain functional connectivity in the low-beta band.

**Interpretation:** The association found between growth measures and brain functional connectivity may reflect a broad deleterious effect of malnutrition on children’s brain development. The mediation effect of functional connectivity on the relation between physical growth and later IQ scores provides the first experimental evidence that brain functional connectivity may mediate the effect of biological adversity on cognitive development.

**Funding:** Bill and Melinda Gates Foundation **(OPP1111625)**

## Introduction

Early exposure to adverse events in childhood has been shown to exert both proximal and distal effects on physical and psychological health and development. Child growth faltering, such as stunting, underweight, and wasting, has been regarded as a primary indicator of malnutrition that affects a large number of children worldwide, especially in low-income countries. According to recent reports by UNICEF, WHO, and the World Bank Group, 159 million children under age five years were classified as stunted, and 50 million were classified as wasted. In the context of the current work, growth faltering has deleterious effects on children’s brain and development cognitive functions^1^,which in turn will greatly impact the degree to which children live up to their developmental potential^2^. Although it is well-known that sufficient nutrients are necessary for brain and cognitive development^3^, the mechanistic pathways by which growth faltering in early childhood affects future cognitive outcomes remain unclear. This is, in part, attibutable to a scarcity of research that has used both neuroimaging and sensitive behaivoral techniques in low-income settings where growth faltering is most prevalent.

In the current study we investigate the consequences of growth faltering on future cognitive and neural outcomes in Bangladeshi children living in impoverished neighborhoods. Specifically, we explore associations between growth faltering and source-space EEG functional connectivity, and whether functional connectivity serves as a mediator of the effects of growth faltering in early childhood on cognitive outcomes. Functional connectivity between EEG oscillatory signals in different frequency bands has been shown to be a useful tool to investigate the development of the efficiency and organization of brain networks^4, 5^. The variation in functional connectivity cannot only be attibuted to changes in brain functioning but may refect anatomical changes in the brain, such as synaptic pruning^6^ and growth of fiber tracts^7^. Abnormal patterns of functional connectivity within certain circuits due to early biological adversity have been linked to deficits in later cognitive performance^8^. It is plausible that communication between cortical areas through neural oscillations in different frequency bands is one of the pathways disrupted by early growth faltering which, in turn, contributes to deficits in cognitive outcomes.

There is a large body of evidence supporting the associations among growth faltering, nutritional deficiencies, and derailed brain development, originating from both human and animal studies^1^. For example, three- to four-month-old infants suffering from malnutrition – indicated by low weight-for-age – showed reduced dendrite growth compared to well-nourished infants^9^. In addition, adults exposed to prenatal famine have white matter (WM) hyperintensities on structural MRI, and it has been suggested that such increases in WM volume may be caused by an inadequate supply of nutrients early in life to sustain and replace catabolized myelin and gliosis after myelin loss^10^. In line with histological evidence from malnourished children, studies of rodents have found that undernutrition is associated with fewer synapses, reduced neuron proliferation, and changes in synaptic structures and pruning^11-13^.

The effects of growth faltering and malnutrition on brain development may lead to poor cognitive outcomes among growth faltered children. A growing literature has demonstrated adverse effects of child growth faltering on children’s cognitive development that, in turn, are believed to contribute to worse educational and labor-market outcomes, including lower incomes and poorer productivity^14, 15^. In addition, stunted growth during infancy and childhood compared to later adolescence is more likely to cause negative long-term effects on adult health and capital^15, 16^. This may be because the first few postnatal years represent a period of rapid neural change^17^ and a critical windows during which experiences have strong effects on brain and cognitive development^18^. Children living in low-resource settings are likely exposed to a variety of biological, psychosocial, and environmental adversities from early in life^19^. Given there is a critical knowledge gap concerning the neural pathways by which growth faltering in early childhood affects cognitive outcomes, it is important to examine the associations among growth faltering, brain functioning, and cognitive outcomes in one study with children living in low-resource settings.

The current study recruited samples of 6- and 36-month-old children residing in an urban slum in Dhaka, Bangladesh. Concurrent growth measures and EEG data were collected at 6 or 36 months. Functional brain connectivity between cortical regions was estimated after cortical source reconstruction of the scalp-level EEG. We focused on the theta, alpha, and low-beta frequency bands because they have been used to examine various cognitive domains such as attention, memory, and emotion processing in infants and young children^20^. Prospective cognitive outcomes were assesed at 27 months for the 6-month-old cohort, and at 48 months for the 36-month-old cohort.

Longitudinal path analysis was conducted to explore: (1) whether growth measures at 6 and 36 months were predictive of future cognitive outcomes; (2) associations between growth measures and brain functional connectivity; and (3) whether brain functional connectivity mediated the effects of child growth on cognitive outcomes. In this model, poverty and family caregiving were statistically controlled as they may confound the relation between growth faltering and neurocognitive outcomes^18, 21^. We hypothesized that child growth faltering would be prospectively associated with worse cognitive outcome in both the 6- and 36-month cohorts. We further hypothesized that brain functional connectivty between multiple regions and circuits in the brain (i.e., whole-brain functional connectivity) would mediate the link between growth and cognitive outcomes.

## Method

### Participants and Growth Measures

The final infant sample consisted of 85 (39M / 46F) children who had usable EEG and growth data at 6 months (M = 6.09, SD = .13). The final toddler sample consisted of 115 (63M / 52F) children who had usable EEG and growth data at 36 months (M = 36.88, SD = .19). Participants’ height-for-age Z-score (HAZ), weight-for-age Z-score (WAZ), and weight-for-height Z-score (WHZ) were calculated based on World Health Organization (WHO) standands. Stunting, underweight, and wasting were respectively defined as the HAZ, WAZ, and WHZ that were <2 SD below the mean of the WHO reference.

### Cognitive Assessment

The cognitive outcome of the infant cohort was assessed with the Mullen Scales of Early Learning (MSEL) at 27 months (M = 26.84, SD = 2.41) for 75 out of the original 85 infants. The scores for four subscales (fine motor, visual reception, receptive language, and expressive language) were standardized and used to calculate a composite T-score reflecting global cognitive development.

The cognitive outcome of the toddler cohort was assessed with the Wechesler Preschool and Primary Scalre of Intelligence (WPPSI-III) at 48 months (M = 48.46, SD = .20) for 109 out of the original 115 children, as children older than 3 years of age tended to demonstrate a ceiling effect on MSEL. The full-scale Intelligence Quotient (IQ) score, a reliable and representative measure of general intellectual functioning, was calculated. The MSEL and WPPSI were both administered by local research assistants and psychologists.

### EEG Data Collection and Processing

EEG was recorded from a 128-channel HydroCel Geodesic Sensor Net (HGSN) that was connected to a NetAmps 300 amplifier (Electrical Geodesic Inc., Eugene, OR) while children watched a screensaver with abstract shapes and soothing sounds for 2 mins. EEG recordings were filtered with a Butterworth band-pass (1 – 25 Hz) filter. The filtered data were then segmented into 1s epochs and inspected for artifacts using absolute and stepwise algorithms, as well as independent component analysis for removing components related to eye movements, blinks, and focal activity (see Supplemental Information for details).

### EEG Functional Connectivity Analysis in the Source Space

The pipeline for the source-space functional connectivity analysis used in the current study is illustrated in Supplemental Figure 1 (also see^22^). Cortical source reconstruction was conducted for the scalp EEG data using realistic head models created for both 6- and 36-month-old cohorts using age-appropriate average MRI templates^23^. Distributed source reconstruction of the EEG time-series was conducted, and reconstructed source activities were segmented into 48 cortical regions of interest (ROIs) using the LPBA40 brain atlas^24^. Functional connectivity between the 48 ROIs was estimated using the weighted phase lag index (wPLI^25^) for different age-appropriate frequency bands: theta (6 mos: 3-6 Hz; 36 mos: 4-7 Hz), alpha (6 mos: 6- 9 Hz; 36 mos: 7-11 Hz), and low-beta (6 mos: 9-15 Hz; 36 mos: 11-15 Hz). Detailed descriptions of the procedures of functional connectivity analysis are included in the Supplemental Information.

**Figure 1.**
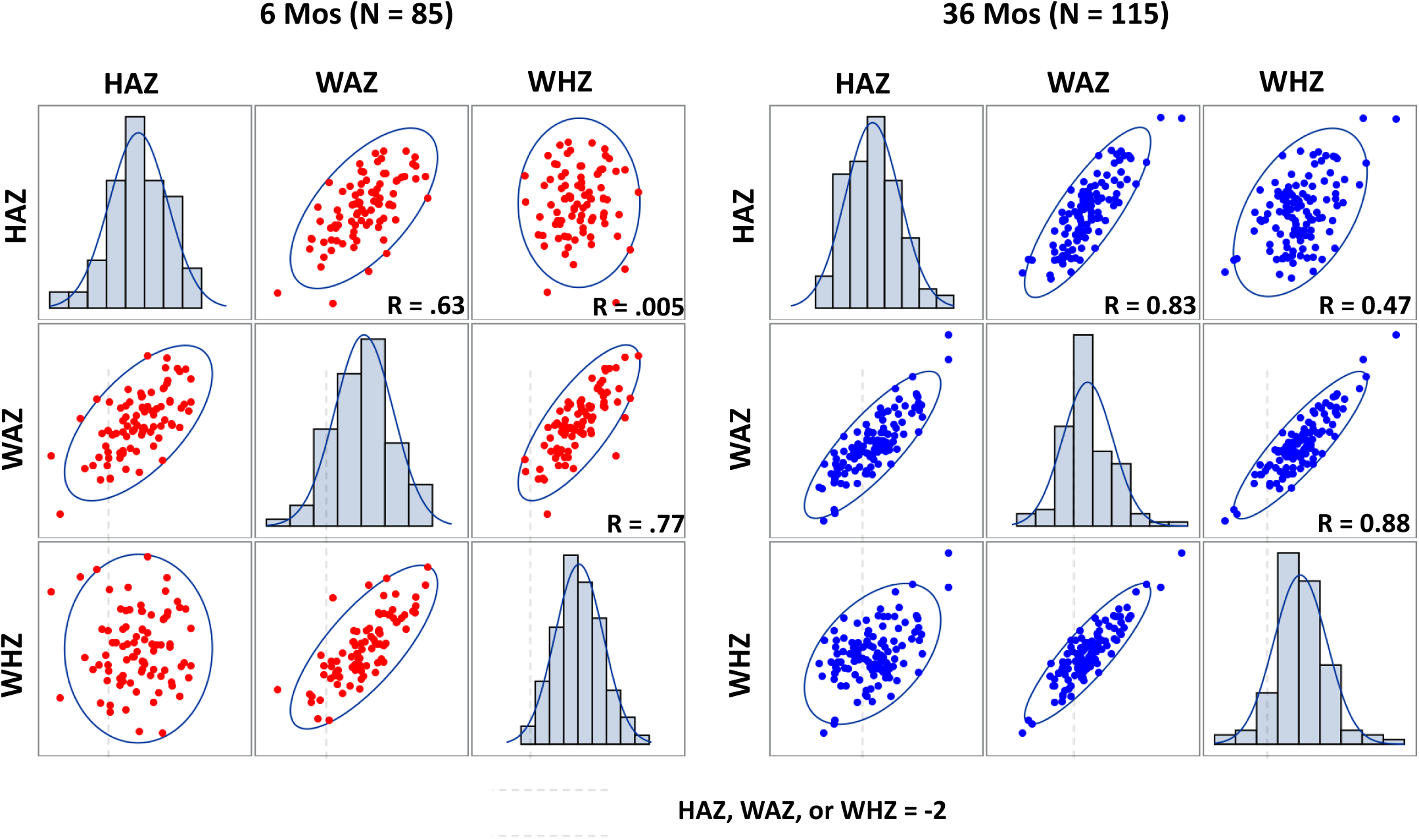
Correlations between the three growth measures and their distribution. The left and panels display the results for the 6- and 36-month-old cohorts respectively.

### Covariates

Poverty was assessed shortly after birth via home assessments and standardized questionnaires. Poverty was defined as a latent factor based on multiple highly-correlated indicators including income-to-needs quartiles, house construction materials, and family assets. Family caregiving activities were assessed at the time of the EEG assessment via maternal interviews with the Family Care Indicators (FCI), which assesses the number of stimulating activities that the parents or other caregivers engaged in with the child within the last 30 days.

### Statistical Analysis

Descriptive statistics, including correlations and partial correlations between variables, were conducted in SAS (version 9.4). The hypothesized mediation model (Figure 5a–c) was tested using longitudinal path analysis in Mplus (version 7.4). This path analysis estimated relations between growth measures and brain functional connectivity, as well as brain functional connectivity and later cognitive outcomes. We explored both direct and indirect effects (via brain connectivity) of growth measures on cognitive outcomes. Missing values for WPPSI scores (N = 6) for the 36-month-old cohort were handled using full-information maximum likelihood (FIML) estimation with robust standard errors. Model fit was evaluated based a non-significant *X*^2^ (p > 0.05), CFI > 0.95, SRMR < 0.08, and RMSEA < 0.06. Indirect effects were estimated using bootstrapping across 10,000 draws with bias-corrected confidence intervals.

## Results

### Growth Measures and Later Cognitive Outcomes

The distribution and correlations of the growth measures for the two cohorts are shown in Figure 1. The average HAZ, WAZ, and WHZ scores for the 6-month cohort were −1.09 (SD = 0.93), −0.71 (SD = 0.98), and 0.047 (SD = 1.02), respectively. The prevalence of stunting was 17.65% (15/85) and higher than that of underweight (9/85; 10.59%) and wasting (2/85; 2.35%). At 6 months, 20 of 85 (23.83%) infants met at least one criterion for poor growth. The average HAZ, WAZ, and WHZ scores for the 36-month cohort were -1.62 (SD=0.95), -1.41 (SD = 1.06), and -0.75 (SD = 1.01), respectively. The prevalence of stunting (40/115; 34.78%), underweight (31/115; 26.96%) and wasting (9/115; 7.83%) at 36 months was much higher that at 6 months. In addition, 47/115 (40.87%) children at 36 months met at least one criterion for poor growth, 24/115 (20.87%) children met criteria for both stunting and underweight, and all 9 wasted children also met criteria for underweight. This result is consistent with the high correlations found between HAZ and WAZ, and between WAZ and WHZ for the 36-month cohort (Figure 1).

Correlations and partial correlations (controlling poverty and family caregiving) between growth measures and cognitive outcomes were analyzed for the two cohorts. There was no correlation between growth measures at 6 months and MSEL scores at 27 months (Table 1a). Positive correlations were observed between the three growth measures at 36 months and IQ at 48 months in the older cohort: HAZ (r = .36, p = .0001; partial r = .21, p = .030), WAZ (r = .39, p < .0001; partial r = .27, p = .0058), and WHZ (r = .31, p = .001; partial r = .23, p = .016) (Table 1b). The growth measures were also analyzed as categorical variables by splitting the 36-month-old children into growth faltered (e.g., stunted group with HAZ < -2) and non-faltered (e.g., middle and high HAZ groups) groups. Growth faltered children were found to have significantly lower IQ scores (Figure 2).

**Figure 2.**
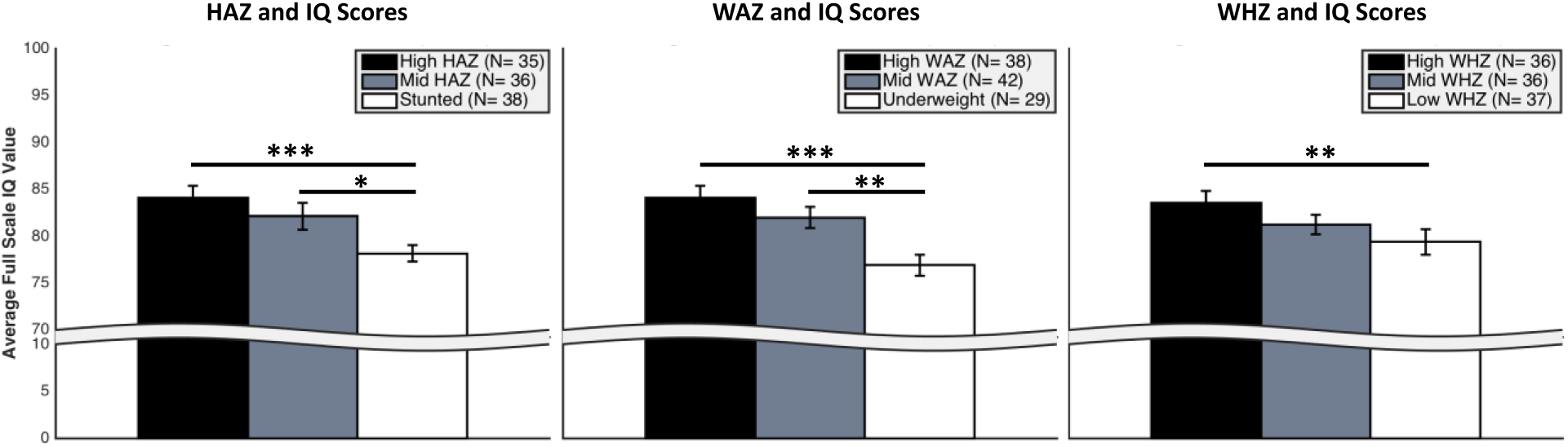
36-month-old children in growth faltered and non-faltered groups showed different IQ scores at 48 months. Black, gray, and white bars represent the IQ scores for the high z-score, middle **z**-score, and faltered (stunted or underweight) growth groups respectively. There were only 9 wasted children, so the WHZ scores were equally divided into three levels to obtain similar observations per WHZ group. Error bars stand for standard errors. * *p* <.05, ** *p* <.0.1, *** *p* <.001.

**Figure 3.**
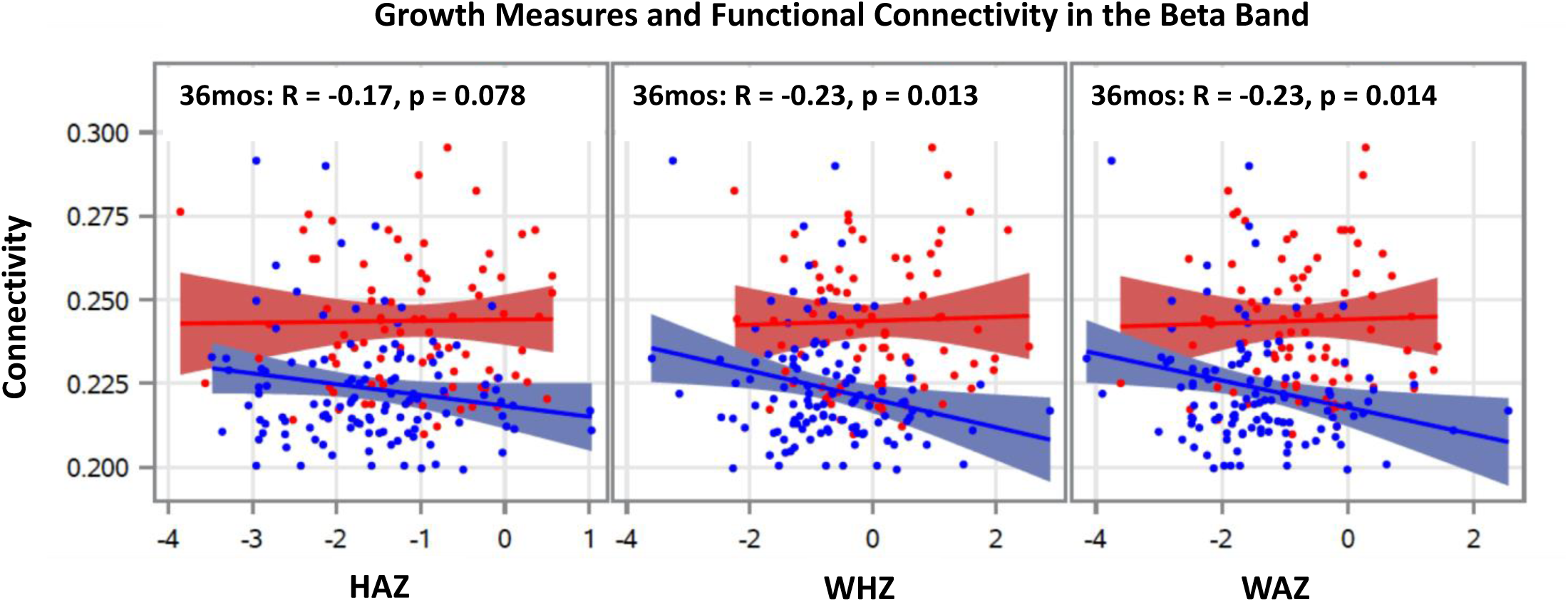
Correlations between growth measures and functional connecitivity in the low-beta bands for the 6- (red) and 36- (blue) month-old cohorts.

**Table 1a.**
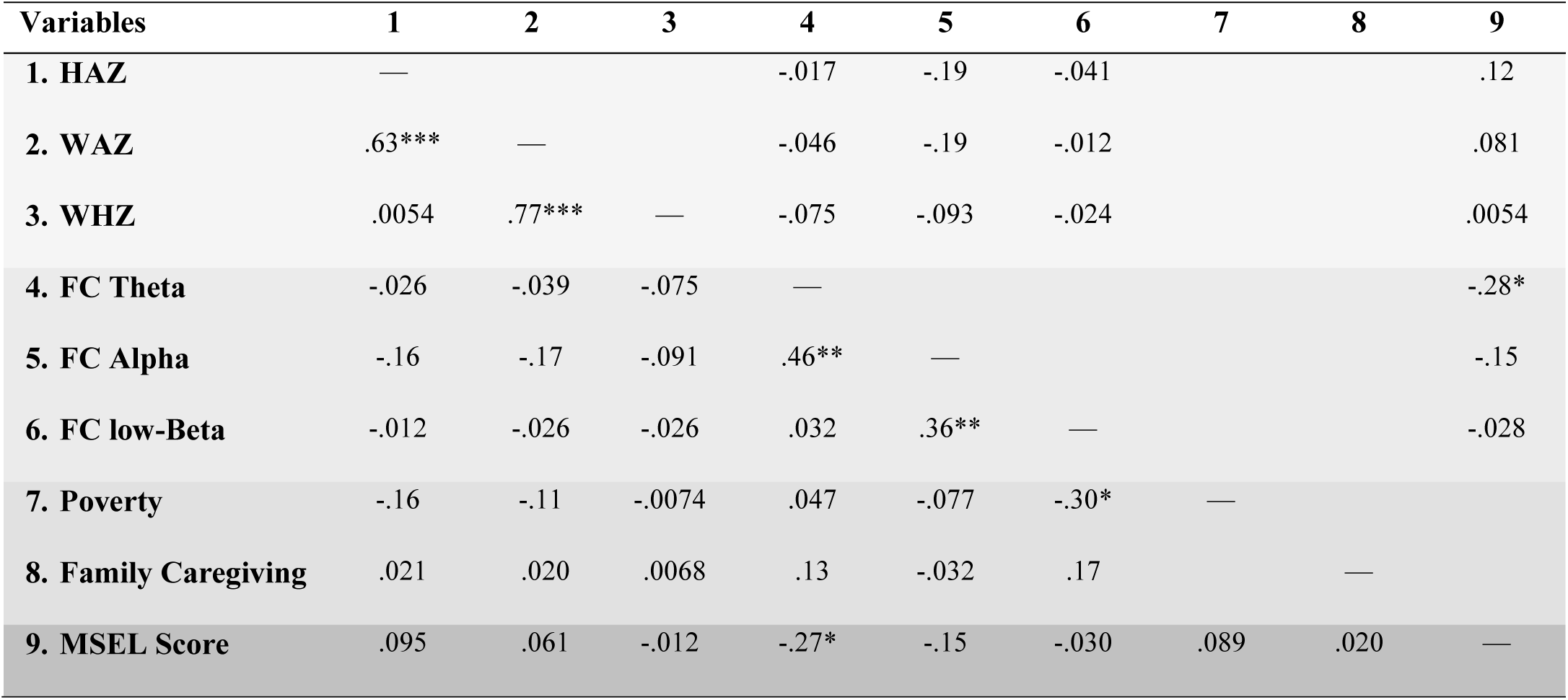
Correlations (bottom half) and Partial Correlations Adjusted for Poverty and Family Caregiving (top half) between the variables for the 6-month-old cohort. *p < .05, **p < .01, ***p < .001.

**Table 1b.** Correlations (bottom half) and Partial Correlations Adjusted for Poverty and Family Caregiving (top half) between the variables for the 36-month-old cohort (*N* = *115*). ^¶^*p* < .08, *p < .05, **p < .01, ***p < .001.

| Variables | 1 | 2 | 3 | 4 | 5 | 6 | 7 | 8 | 9 |
| --- | --- | --- | --- | --- | --- | --- | --- | --- | --- |
| **1. HAZ** | — |  |  | -.25** | -.15 | -.25** |  |  | .21* |
| **2. WAZ** | .83*** | — |  | -.32*** | -.19* | -.28** |  |  | .27** |
| **3. WHZ** | .48*** | .88*** | — | -.28** | -.17† | -.28** |  |  | .23* |
| **4. FC Theta** | -.16† | -.18† | -.15 | — |  |  |  |  | -.10 |
| **5. FC Alpha** | -.12 | -.16 | -.15 | .53*** | — |  |  |  | -.21* |
| **6. FC low-Beta** | -.16† | -.23* | -.23* | .35*** | .56*** | — |  |  | -.26** |
| **7. Poverty** | -.49*** | -.46*** | -.33*** | -.11 | -0.011 | -.10 | — |  |  |
| **8. Family Caregiving** | .13 | .10 | .027 | -.047 | -.053 | .12 |  | — |  |
| **9. IQ Score** | .36*** | .39*** | .31** | -.05 | -.19* | -.18† | -.39*** | .22* | — |

### Growth Measures and Brain Functional Connectivity

There was no correlation between growth measures and functional connectivity for the 6-month cohort. In contrast, growth measures were negatively correlated with brain functional connectivity in the low-beta band for the 36-month cohort: HAZ (r = -.16, p = .078; partial r = -.25, p = .0072), WAZ (r = -.23, p = .013; r = -.28, p = .0026), and WHZ (r = -.23, p = .014; partial r = -.28; p = .0030) (Table 1). Strong partial correlations between growth measures and brain functional connectivity in the theta band were also observed for the 36-month cohort (Table 1b). As above, these children were then divided into growth faltered and non-faltered groups. Brain functional connectivity in the low-beta band was found to be higher and with stronger connections between brain regions for the faltered than non-faltered groups (Figure 4). No significant correlations or partial correlations were found for the alpha band.

**Figure 4.**
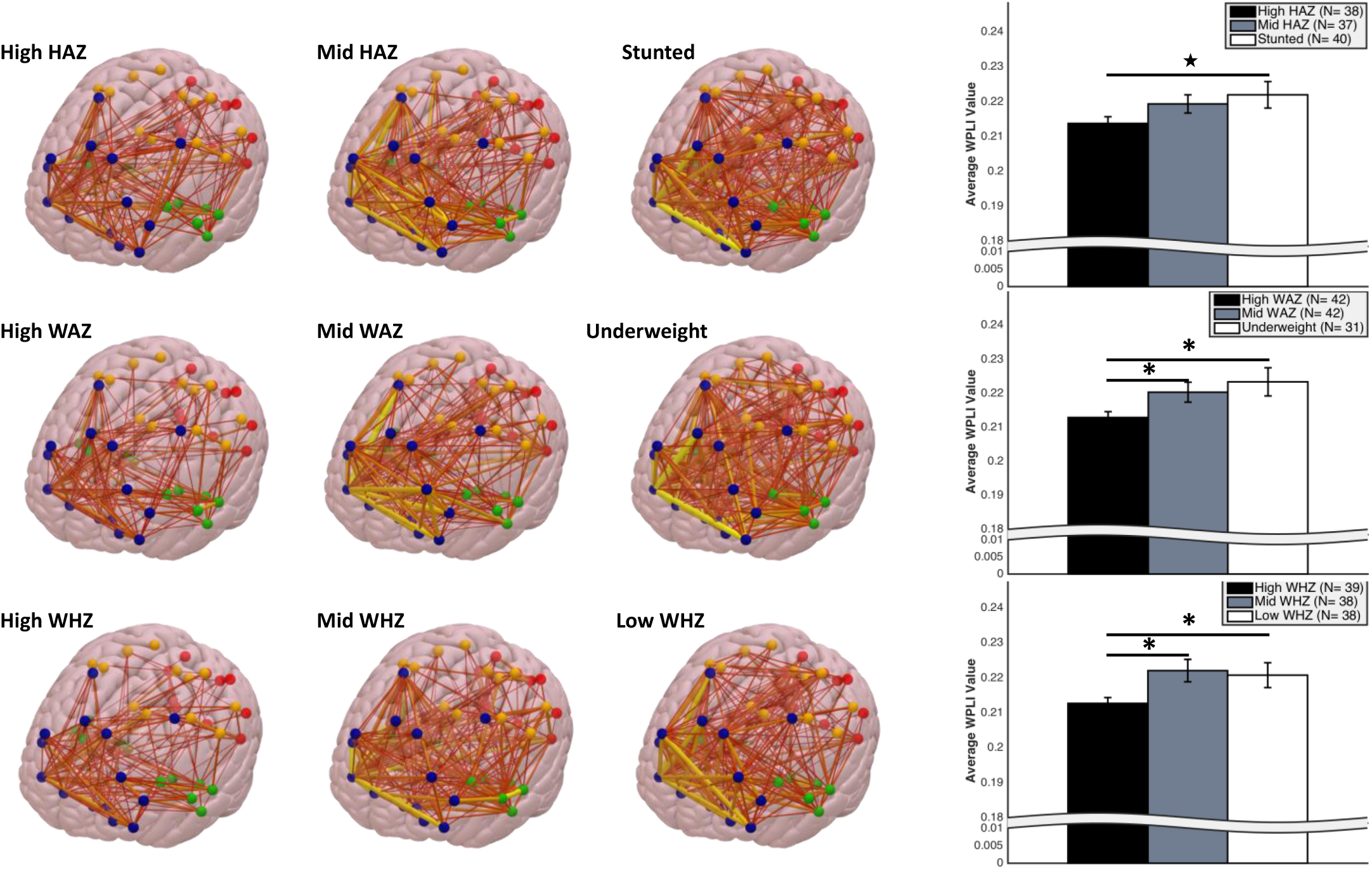
36-month-old children were divided into growth faltered and non-faltered groups and brain functional connectivity in the low beta band is illustrated for each group in the brain (left side), as well as by bars (right side). Black, gray, and white bars represent the IQ scores for the high z-score, middle z-score, and faltered (stunted or underweight) growth groups respectively. There were only 9 wasted children, so the WHZ scores were equally divided into three levels to obtain similar observations per WHZ group. Error bars stand for standard errors. * *p* <.05,★ *p* < .07.

### Brain Functional Connectivity and Later Cognitive Outcomes

A negative correlation was found between brain functional connectivity in the theta band at 6 months and MSEL score at 27 months in the infant cohort, r = -.27, p = .026; partial r = -.28, p = .022. Negative correlations were also found between brain functional connectivity in the alpha (r = -.19, p = .044; partial r = -.21, p = .028) and low-beta (r = -.18, p = .053; partial r = -0.25, p = .0076) bands at 36 months and IQ scores at 48 months in the toddler cohort.

### Mediation Effect of Brain Functional Connectivity

Mediation analysis was not run for the 6-month infant cohort given the absence of correlations between growth measures and cognitive outcomes, and between growth measures and brain functional connectivity. Mediation models were tested for the 36-month cohort, with the association between growth measures (HAZ, WAZ, and WHZ) and future IQ hypothesized to operate through brain functional connectivity in the low-beta band. Model fit is reported in the Supplemental Information. The mediation models revealed that each growth measure was associated with brain functional connectivity at 36 months, which in turn predicted IQ at 48 months (Figure 5a-c). We observed indirect effects of growth measures on IQ through brain functional connectivity in the low-beta band in all three models. The presence of such indirect effects suggests that the effect of growth on IQ was mediated by brain functional connectivity.

**Figure 5.**
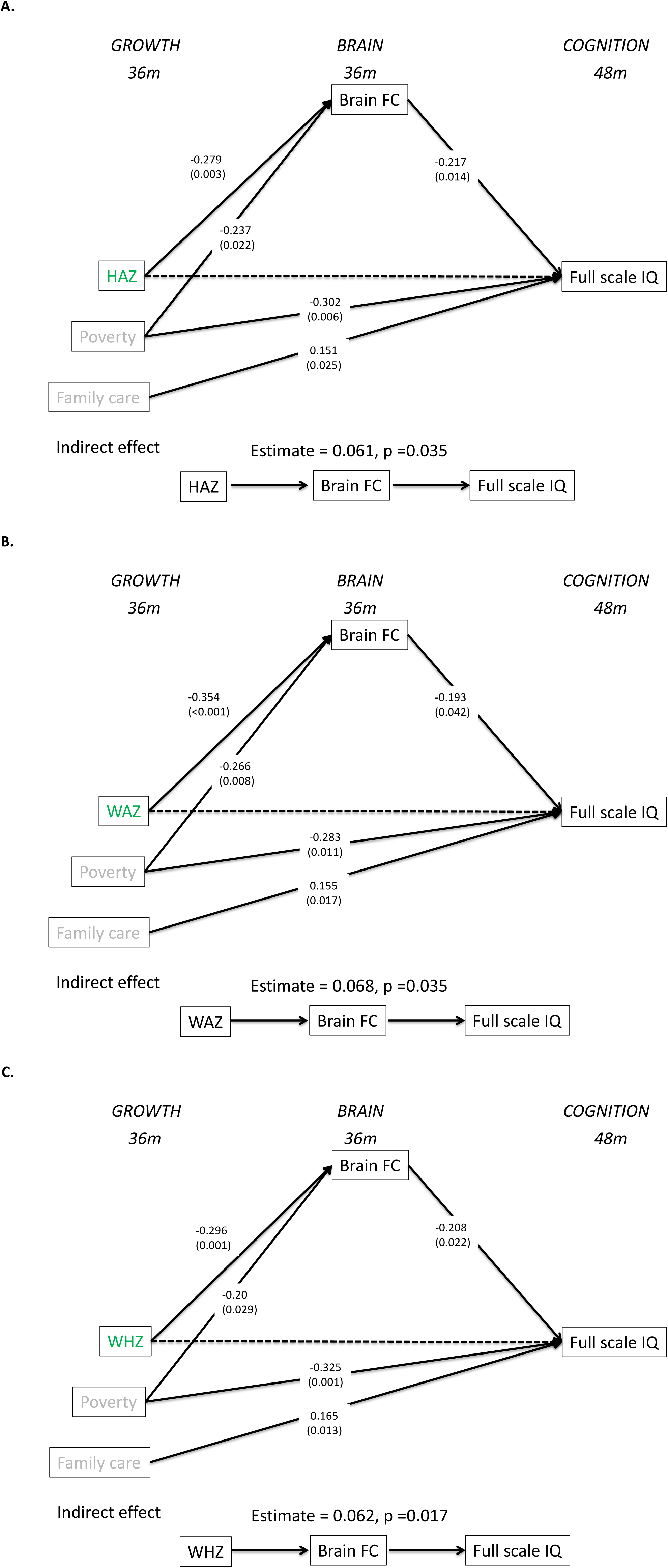
Multivariate mediation Models tested with longitudinal path analysis a) the model with HAZ, b) the model with WAZ, and c) the model with WHZ. Variables colored in grey are covarites. The numbers in parentheses are the p-values, and the numbers above the p-values are the estimates.

## Discussion

Here for the first time we demonstrate that functional brain connectivity in theta and low-beta bands, respectively, predicted future cognitive performance in the 6- and 36-month cohorts and that functional brain connectivity in the low-beta band at 36 months mediated the effect of growth faltering on IQ one year later.

The high prevalence of growth faltering in the 36-month cohort highlights the severity of deficits in neurobiological development in this sample of Bangladeshi children. In this cohort, ~35%, 27% and 8% of the children were classified as stunted, underweight, or wasted (respectively), and approximately 40% met at least one criterion of poor physical growth. These prevalence rates are comparable to a report by the Bangladesh Demographic and Health Servey in 2014, which showed that the prevelance of stunting, underweight, and wasting is 33.5%, 33%, and 14%, respectively, among children under age 15. The prevalence of growth faltering at 6 months was relatively lower than that of the 36-month cohort, with approximately 25% of the infants meeting criteria for stunting, underweight, or wasting. The dramatic difference in the prevalence of growth faltering between the 6- and 36-month cohorts highlights the importance of continuing to monitor the growth trajectories of children in low-resource settings, and the need to continually foster growth, over the first three years of life.

The effect of growth faltering at 36 months on children’s IQ scores at 48 months supports the notion that exposure to early adversity increases the risk of undermining healthy cognitive development. This finding is consistent with previous reports of associations between growth stunting and performance on cognitive tests^26-28^. Since stunting, underweight, and wasting are frequently used as indicators of malnutrition, the difference in IQ scores found between growth faltered and non-faltered groups highlights the importance of adequate nutrients for normal cognitive development and suggests that interventions to facilitate physical growth may promote gains in cognitive performance among those who experience growth faltering.^29^ Consistent with the literature suggesting the robust role of poverty in cognitive development^21, 30^, poverty was also found to explain a large amount of variance in IQ scores. Given the large number of variables that covary with poverty, future research should be mindful that deficits in cognitive development are likely caused by a constellation of interacting factors for children living in low-income countries^31^.

The correlations found between growth measures and whole-brain functional connectivity for the 36-month cohort may reflect a broad deleterious effect of growth faltering on children’s brain functioning. The neural oscillations in the theta and low-beta bands have been associated with cognitive functions such as sustained attention, executive attention, and working memory^32-35^. Atypical patterns of functional connectivity in these frequency bands may therefore be associated with deficits in cognitive processes in a number of domains. The absence of an association between growth measures and functional connectivity in the alpha band could be due to the fact that synchronization in this band was suppressed during the experiment when children were watching (attending to) a screensaver with abstract shapes, as alpha is known to be attenuated whilst engaging in attention-demanding tasks.

Although a significant association between growth faltering and brain functional connectivity was observed, the direction of this association was contrary to our hypothesis. That is, we observed that growth faltering was associated with higher functional connectivity. One explanation for this finding is that synaptic pruning among growth faltered children may be delayed due to malnutrition, in turn leading to a proliferation of connections^10^, as well as less organized and more superfluous pathways between brain networks. Children’s experience plays a key role in synaptic pruning, which begins in the first year after birth and continues through adolescence^36^. Delayed synaptic pruning or accelerated proliferation may occur in stunted children due to a lack of stimulation and input from the environment. A second explanation is that higher connectivity reflects an adaptive or compensatory neural response to delayed brain anatomical development in growth faltered children. For instance, higher connectivity could indicate less efficiency of neural communications between cortical regions which requires more effort for children who are stunted, underweight, or wasted.

A particularly compelling finding of the current study is that the effect of growth faltering on children’s IQ operated through brain functional connectivity in the low-beta band. Biological adversity related to inflammation has recently been shown to impact functional connectivity and cognitive abilities by studies using fMRI techniques^8, 37^. Our finding adds to this growing literature by showing that functional connectivity mediates the relationship between biological adversity and later cognition, which means functional connectivity may serve as a neural pathway by which exposure to early adversity affects cognitive outcomes. As a practical matter, the current study suggests that EEG – a measure that is less expensive and easier to implement than MRI – also has the sensitivity to detect differences in brain functional connectivity in relation to early adversity in low resource settings.

The absence of an association of growth faltering to functional connectivity and cognitive outcomes in the 6-month cohort was inconsistent with the study hypothesis. The impact of growth faltering on brain and cognitive functioning may require a greater degree of maturation before its effects fully manifest, which suggests the need for future research following these children longitudinally throughout childhood. In contrast, brain functional connectivity at 6 months is already predictive of cognition at 27 months, suggesting that measurement of neural function during infancy may be one means of capturing risk for future cognitive difficulties among children in low-resource settings.

## Conclusion

The current study is the first attempt to investigate the associations among growth faltering, brain functional connectivity, and cognitive outcomes in children living in low-income countries. Our findings indicate that growth faltering, an indicator of malnutrition, is associated with exaggerated functional connections in the brain, which predicts poor cognitive outcomes later in life. These findings advance our understanding of the neural pathways by which faltered growth could influence cognitive development, and this advancement may have a substantial impact on developing efficient interventions for children living in low-income countries.

